# Towards a unifying diversity-area relationship (DAR) of species- and gene-diversity

**DOI:** 10.1101/2020.05.16.099861

**Authors:** Zhanshan (Sam) Ma, Aaron M. Ellison

## Abstract

**Aim:** The microbiome as a biogeographic entity can be investigated, to the minimum, from two perspectives: one is the spatial/temporal distribution of species (or any level of the operational taxonomic unit or OTU) diversity, and another is the spatial/temporal distribution of metagenomic gene diversity. Both are necessary for comprehensive understanding of the taxonomical, ecological, evolutionary and functional aspects of the microbiome biogeography. Here we propose to investigate the metagenomic diversity-area relationship (*m*-DAR), which is a transformation of the species-DAR (*s*-DAR) that extended the classic SAR (species-area relationship) by replacing the species richness with general species diversity measured in Hill numbers.

**Innovation:** The *m*-DAR and *s*-DAR, using the same mathematical models, offer a unifying tool for investigating the biogeography of microbiome from ecological, metagenomic and functional perspectives. Specifically, we investigate *m*-DAR of the human gut metagenome in terms of the MG (metagenomic gene) and MFGC (metagenome functional gene cluster) respectively, by sketching out the DAR-profile, PDO (pair-wise diversity overlap) profile, MAD (maximal accrual diversity) profile, and RIP (ratio of individual- to population-diversity) profile at each scale. These profiles constitute our unifying DAR toolset and can be applied to any microbiomes beyond the human gut microbiome.

**Main conclusions:** We demonstrate the construction and applications of the *m*-DAR and its associated four profiles with six large datasets of the human gut metagenomes including three microbiome-associated diseases (obesity, diabetes, IBD) and their healthy controls, supported with randomization tests to determine the differences between healthy and diseased treatments in their *m*-DAT parameters. Theoretically, our study presents a successful case to demonstrate the feasibility of unifying systematic biogeography *vs*. evolutionary biogeography, of an inclusive biogeography of plants, animal and microbes. Practically, our approach offers an important tool for investigating the spatial scaling of human metagenome diversity in a population (cohort) and its relationship with individual-level diversity.

## Introduction

The term *biogeography* was coined by Clinton Hart Merriam (1855-1942) (Cited in Ebach 2007), and a number of definitions and interpretations have been attached to it, each with its own flavor toward a particular field of study (Ebach 2007). For example, microbial biogeography has been defined as the study of spatial and temporal distribution of microbial diversity (Martiny *et al*. 2006, Hanson et al. 2012, Costello et al. 2012, van der Gast 2013, 2015), while the molecular ecologist may define biogeography to be the study of gene lineage. As argued by Ebach (2007), whom we agree with, biogeography is suffering form something of an identity crisis when the term is used as a ‘catch-all’ for characterizing the spatial aspect of a particular life-science field, rather than being treated as an independent field of science. In the present study, we are interested in the question: should microbial biogeography also be concerned with the spatial or temporal distribution of the gene or metagenome (*i.e*., the total genomes of all member species in a microbial community or microbiota)? We answer the question by performing a case study— extending the SAR (species area relationship), one of a few classic laws in ecology and biogeography and also one of the most general yet protean patterns in nature (Lomolino 2010, Whittaker & Triantis 2012), to the microbial genes. Specifically, we transform and expand the DAR (diversity-area relationship) of microbiome species or OTUs (operational taxonomic units) (Ma 2018a, 2018b), *i.e*., *s*-DAR or simply DAR hereafter, which extended the classic SAR to general diversity-area scaling, into metagenome-DAR (metagenome-diversity area relationship) or *m*-DAR.

The documented studies of the classic SAR pattern can be traced back to as early as the 19^th^ century (Watson 1835), and other early pioneering and seminal studies such as Arrhenius (1921), Preston (1960, 1962), MacArthur & Wilson (1967), Connor & McCoy (1979), Rosenzweig (1995) played an important role for the establishment of island biogeography and biogeography at large. For example, the earliest documentation of SAR was by Watson (1835), whose work was recognized to even have a significance influence on Darwin’s (1859) development of evolutionary theory because both evolution and ecology have to deal with the distribution of biodiversity on the earth planet. The difference between ecological and evolutionary perspective is simply the *time* scale.

One of several internal divisions of biogeography has been the distinction between *systematic biogeography* vs. *evolutionary biogeography*. Outside microbial biogeography, it has been argued that the differences between the two biogeographies reflected the schisms between evolutionary biology and systematics. Furthermore, the continuing conflict is not due to personality, methodology, or implementation, and instead, is due to opposing *intent* (Ebach 2007). One intention is to use trusted models to generate all possible historical scenarios by opting for a unifying approach. Alternatively, one may intend to classify and identify all possible biota and form a classification that should exhibit universal patterns (Ebach 2007). There is a need to bridge both approaches (fields) because as Ebach (2007) stated: “In fact, the two biogeographies are equally ecological, historical, and paleontological” (also see Riddle and Hafner 2007). We hope our work is a useful contribution to bridge the gap. According to this classification, SAR or DAR should belong to the former branch and the *m*-DAR to the latter. If our transformation from DAR to *m*-DAR is successful, we suggest that a unifying biogeographic approach is feasible. We demonstrate the transformation and feasibility with extensive datasets from the human microbiome project (HMP), specifically 6 datasets of the human gut metagenome including three high-profile microbiome-associated diseases (obesity, diabetes, and IBD [inflammatory bowel disease]) and their healthy controls.

Recent advances in human microbiome research have revealed that human gut microbiome is highly personalized (heterogeneous), which means that, in general, neither the distribution of microbial species (ecological) diversity nor the distribution of metagenome (total genomes of microbes a microbiome carries) diversity within a human population is homogenous. For example, if each individual has the exactly same number of metagenomic genes (MG), then the population should have the same number of MGs as each individual. In reality, the number of gut MGs an individual carries varies hugely from individual to individual and from time to time, which makes the calculation of metagenomic gene numbers (*i.e*., gene richness) in a population rather complex. In other words, unlike the *fixed* number of genes a human genome contains (approximately 20,000), the number of MGs within a metagenome of a human microbiome contains is highly variable. In contrast, the *standard error* of the MG in a human cohort is close to the number of human genes (*see* Table 1). One component of our *m*-DAR analysis, the RIP (the ratio of individual- to population-diversity) profile readily resolves the above issue.

**Table 1.** Brief information on the metagenome datasets used to demonstrate *m*-DAR models

| Disease | Treatments | Number of Samples | Number of samples selected for *m*-DAR | Mean number of genes & Std. Err | Mean number of MFGC & Std. Err (eggNOG) | Reference |
| --- | --- | --- | --- | --- | --- | --- |
| Obesity | Lean | 96 | 96 | 973, 838±26, 969 | 245 ± 2 | Qin *et al.* 2010<br>Chatelier *et al.* 2013 |
|  | Obesity | 17 | 0 (not used) |  |  |  |
|  | Overweight | 168 | Randomly sampled 96 | 937, 187±24,350 | 239 ± 2 |  |
| Type-II Diabetes | Healthy | 74 | 74 | 523, 793±17, 829 | 223 ± 1 | Qin *et al.* 2012 |
|  | Diseased | 71 | 71 | 504, 712±23, 219 | 222 ± 2 |  |
| IBD | Healthy | 24 | 71 | 962, 413±30, 600 | 245 ± 2 | Nielsen *et al.* 2014 |
|  | Healthy Relative | 47 |  |  |  |  |
|  | CD | 21 | 0 (not used) |  |  |  |
|  | UC | 127 | Randomly sampled 71 | 838, 009±24, 347 | 236 ± 2 |  |

If we further pursue the calculation of *metagenomic gene* (MG) *diversity* (beyond gene richness) of a *population*, which consider both the kinds (or ‘species’) of genes and the abundance of each gene, the problem becomes even more complex, because the diversity is measured in entropy and entropy is a non-linear function of gene abundances (Ma & Li 2018). In this paper, we tackle this problem of metagenome diversity changes (scaling) with the change of human population size. This problem is both theoretically interesting and practically significant. Theoretically, the problem is essentially the counterpart of ecological DAR (Ma 2018a, 2018b), which is a superset of classic SAR in ecology and biogeography, as mentioned previously. Practically, recent studies have found that we are losing biodiversity in arguably the closest ecosystem (*i.e*., the gut microbiome) to our body, most likely, due to the modern life style and the use and occasionally abuse of antibiotics. The loss of biodiversity in gut microbiome is likely related to many of our diseases such as obesity, IBD, diabetes and hence has far reaching implications to our health. For this reason, scientists have been calling for the conservation/restoration of our ‘native’ gut microbiome. Obviously, the conservation/restoration of gut microbiome (metagenome) diversity will need to monitor the change of gut microbiome (metagenome), in both microbial ecological diversity and metagenome diversity of the genes (carried by the microbes). The metagenome-DAR or *m*-DAR (to be developed in this study) and the original ecological or species *s*-DAR (Ma 2018a, 2018b) offer a unifying approach for measuring both microbiome species diversity and metagenome diversity scaling within a human population. Although, we use the datasets of human microbiome to demonstrate the proposed *m*-DAR, the concepts, principles and models should be equally applicable to most other microbiomes on the planet such as plant and ocean microbiomes. An immediate application of our work can be the estimation of the potential (or ‘dark’) diversity, which refers to species that is absent locally but present in regional species pool, across the spectrum of rarity-commonness (Partel *et al*. 2011, Real *et al*. 2017, Ma 2019). For example, how many total metagenomic genes, commonly encountered genes with moderate abundance (frequency), or most frequently encountered (dominant) genes are there in a human population? For another example, the ratio of individual (local) to population (regional) level metagenome diversity can also be estimated from the *m*-DAR scaling parameters. These statistics and parameters can be of significant importance for understanding the biogeography of the human metagenome, and should be equally applicable for other microbiomes.

## Materials and Methods

### The six metagenome datasets of the human gut microbiome

Table 1 below listed the brief information on the metagenome datasets we use to demonstrate the *m*-DAR analysis. To keep the balanced sample sizes between the healthy and diseased treatments (groups), in some cases, we randomly discarded certain number of samples, so that the model parameters can be compared appropriately between the healthy and diseased treatments.

### Definitions and computational procedures for *m*-DAR (metagenome-diversity area relationship)

The process for constructing *m*-DAR models consists of the following three steps (Fig 1): (*i*) Bioinformatics analysis of the metagenomic sequencing raw reads (data) to get MGA (metagenomic gene abundance) tables and MFGC (metagenome functional gene cluster) tables; (ii) Computing metagenome diversity for the MG (metagenomic gene) and MFGC, respectively; (iii) Fitting the *m*-DAR models and construct the four profiles, for MG and MFGC, respectively.

**Fig 1.**
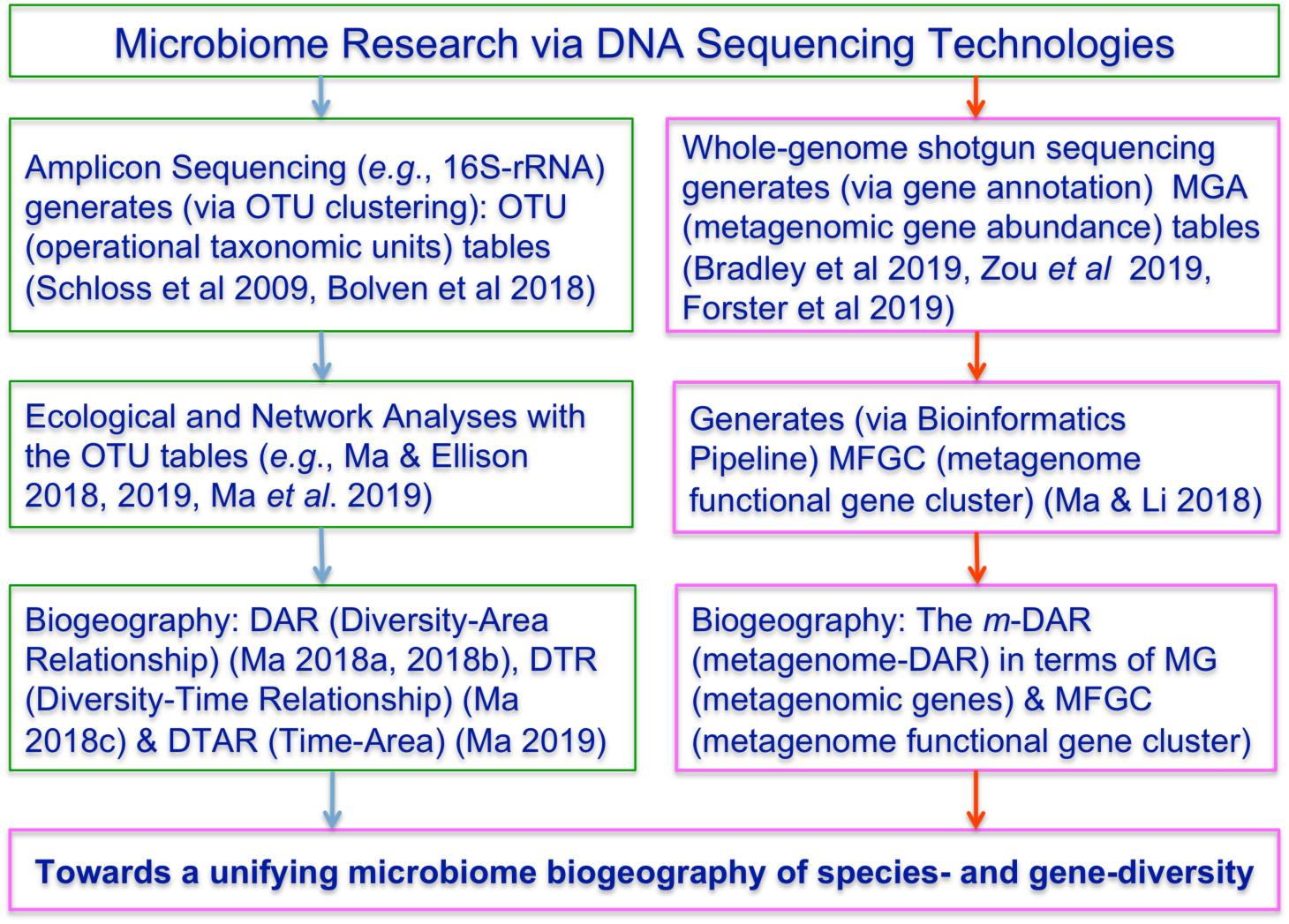
A diagram showing the bioinformatics pipelines and ecological tools to establish a unifying biogeography for microbiome species- and gene-diversity

#### Computing the MGA (metagenomic gene abundance) tables and MFGC (metagenome functional gene cluster abundance) tables

To construct m-DAR models, the first step is to compute the MGA tables from metagenomic sequencing (whole-genome shotgun sequencing) reads by using standard bioinformatics software pipelines (Zhu *et al* 2010, Qin et al. 2010, 2013, Chatelier et al. 2013, Li et al 2014, Xiao et al. 2015, 2016, Wang & Jia 2016, Sczyrba et al. 2017, Ma & Li 2018). The formal definitions for MFGC can be found in Ma & Li (2018). An MFGC is a cluster of metagenomic genes (MG) with the same group of functions. Therefore, a large number of MGs (hundreds of thousands) are usually classified into a small number of MFGCs (hundreds). For example, in Table 1, the number of MGs of the ‘lean’ treatment in the obesity case study was 973,838, but the number of MFGCs was 245 only. Two commonly used databases for identifying the MFGC are eggNOG (protein functions) and KEGG (metabolic pathways) databases, based on which two different kinds of MFGCs can be identified.

#### Computing metagenome diversity in Hill numbers

Similar to the computation of ecological diversity (*aka* species diversity or community diversity, or occasionally microbiome diversity), Ma & Li (2018) defined metagenome diversity for MG and MFGC, respectively in terms of the Hill numbers. Hill numbers are a series of entropies, corresponding to different diversity order (*q*=0, 1, 2,…), defined for the MG (or MFGC) abundance distribution (table) data. The metagenome diversity of MG is measured at single-gene level, and the metagenome diversity of MGFC is measured at functional gene cluster level. Regarding the metagenome diversity of MFGC, two sub-types were distinguished based on how the individual gene abundance information is utilized in defining MFGC, MFGC-I (type-I MFGC) ignores the gene abundance of individual genes and only counts the number of gene kinds (or species) in a cluster, and MFGC-II (type-II MFGC) considers both gene kinds (or species) and abundances (Ma & Li 2018). However, when diversity order *q*=0, this distinction does not make any difference because when *q*=0, the abundance information does not weigh in anyway. As it is explained later, for MFGC, we only build m-DAR models for diversity order *q*=0. Therefore, for the purpose of constructing m-DAR models for MFGC, there is no need to distinguish between the two types of MFGCs, and we just use the term “MFGC” for simplicity in this article. Although distinguishing the two types of MFGC regarding the utilization of the individual abundance information is not necessary as explained here, there is another typing of MFGC based on the functional gene databases (KEGG or eggNOG) used, from which MFGCs were compared and computed, is necessary as explained in the previous subsection. That is, we formally distinguish MFGC as two types, one is based on KEGG database (for metabolic pathways) and another is based on eggNOG (protein functions).

The Hill numbers, originally introduced as an *evenness* index from economics by Hill (1973), was reintroduced into ecology by Jost (2007) and Chao *et al.* (2012, 2014) who further clarified Hill’s numbers for measuring alpha diversity, and can be adapted to measure metagenome diversity of MG or MFGC as:

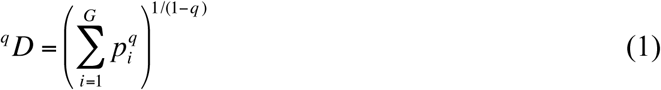

where *G* is the number of MG/MFGC (*kinds* or *species*, equivalent to the number of species for ecological diversity or species diversity), *p*_i_ is the relative abundance of *i*-th MG/MFGC, *q* is the order number of diversity. When *q*=0, the diversity ^*0*^*D*=*G*, which is the number of MGs/MFGCs, and is termed MG/MFGC richness. When *q*=1, ^*1*^*D* is a function of Shannon index, and the diversity is weighted *proportionally* by the gene (MG/MFGC) frequency. When *q*=2, ^*2*^*D* is a function of Simpson index and the diversity is weighted in favor of the dominant (more abundant) genes (MGs/MFGCs) (Ma & Li 2018).

#### Constructing *m*-DAR models

We propose to use the same mathematical functions and computational procedures used for ecological DAR models (Ma 2018a), to construct *metagenomic* DAR (*m*-DAR) models here.

The metagenomic diversity-area relationship (*m*-DAR) is:

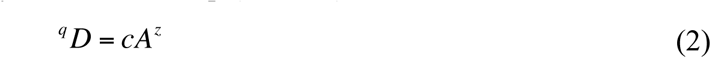

where ^*q*^*D* is the metagenome diversity measured in the Hill numbers of the *q*-*th* order, *A* is *area* (the number of subjects whose metagenome diversity are sampled), and *c* & *z* are parameters. We term this model *m*-DAR power law (PL) model, which has the same mathematical form as the classic SAR and species-DAR (or *s*-DAR, Ma 2018a).

The slope (*z*) of the *m*-DAR PL model (also known as the diversity-scaling parameter) is the ratio of metagenome diversity accrual rate to area increase rate. The relationship between DAR-PL model parameter (*z*) and the diversity order (*q*), or *z-q* trend, was defined as the DAR profile (Ma 2018a). It describes the change of diversity scaling parameter (*z*) with the diversity order (*q*), comprehensively. Similarly, we extend this definition of DAR profile to *m*-DAR profile.

A slightly modified PL model, the power law with exponential cutoff (PLEC) model, originally introduced to SAR modeling by Plotkin et al. (2000) and Ulrich & Buszko (2003), respectively (also see Tjørve 2009), was utilized for DAR modeling (Ma 2018a). Here, we use the PLEC model for *m*-DAR analysis, which is of the form:

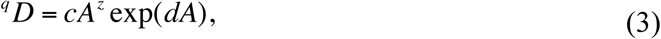

where *d* is a third parameter (tapper-off parameter) and should be negative in DAR scaling models, and exp(*dA*) is the exponential decay term that eventually overwhelms the power function behavior at very large value of *A*. The justification for adding the exponential decay term is because both the human body and metagenomic genes carried by the human microbiome are finite, and there should be a taper-off parameter to reflect the finite size of metagenome diversity.

The following log-linear transformed equations (4)(5) can be used to estimate the model parameters of Eqns. (2)(3), respectively:

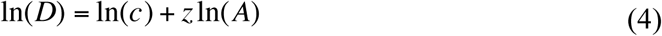

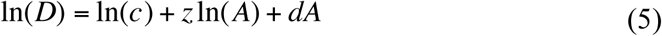

Both linear correlation coefficient (*R*) and *p*-value can be used to judge the goodness of the model fitting. In fact, either of them should be sufficient to judge the suitability of the models to data. Three advantages are associated with the linear-transformed fitting: (*i*) simplicity in computation, (*ii*) parameter z is scale-invariant with eqn. (4), (*iii*) the ecological interpretation of scaling parameter is preserved with eqn. (5). The scaling parameter *z* is also termed the slope of the power-law DAR, because *z* represents the *slope* of the linearized function in log–log space.

There is an artificial but significant difference between the human microbiome and other biomes in nature. That is, there is usually a natural spatial order or arrangement for plants and animals, but there is not such a natural order among humans (the hosts of human microbiomes) because human residences or even the sampling locations are near totally artificial and make little biomedical sense naturally. To deal with the lacking of a definite natural spatial order among individual subjects, we first randomly permutate the orders of the samples of a treatment and obtain the total permutations of the possible orders. We then randomly select 50 (for MG, the consequent computation with MG-DAR is extremely time-consuming) or 100 (for MFGC, the computation with MFGC-DAR is much less time-consuming) orders from the total permutations, and fit the m-DAR model for each of the 50 or 100 orders. Finally, we take the average parameters from the 50 (or 100) permutations and treat the average parameters as the final m-DAR models. Among the 50/100 times of repetition, we eliminate those failed modeling attempts (with *p*>0.05). In PLEC modeling, we also remove the repetitions with *A*_*max*_<0 (which is biologically infeasible). The number of removed repetitions was actually extremely small, as shown in the column of *N*\* for each DAR model (PL or PLEC) in Table 2.

**Table 2.** The parameters of *m*-DAR (metagenome diversity-area relationship) models fitted for metagenomic gene (MG) diversity, averaged from 50 times of re-sampling

| Diversity Order | Study Case | Treatments | m-DAR PL (power law) Model |  |  |  |  |  | m-DAR PLEC (power law with exponential cutoff) Model |  |  |  |  |  |  |  |
| --- | --- | --- | --- | --- | --- | --- | --- | --- | --- | --- | --- | --- | --- | --- | --- | --- |
| | | | $z$ | $\ln(c)$ | $g$ | $R$ | $p$-value | $N^*$ | $z$ | $d$ | $\ln(c)$ | $R$ | $p$-value | $A_{max}$ | $D_{max}$ | $N^*$ |
| $q=0$ | Obesity | Lean | 0.358 | 14.224 | 0.718 | 0.975 | 0.000 | 50 | 0.501 | -0.005 | 13.969 | 0.994 | 0.000 | 94 | 6871100 | 50 |
|  |  | Overweight | 0.367 | 14.146 | 0.711 | 0.975 | 0.000 | 50 | 0.509 | -0.005 | 13.890 | 0.993 | 0.000 | 96 | 6629101 | 50 |
|  | Type-II diabetes | Healthy | 0.435 | 13.503 | 0.648 | 0.982 | 0.000 | 50 | 0.577 | -0.007 | 13.279 | 0.994 | 0.000 | 86 | 4305968 | 50 |
|  |  | Disease | 0.468 | 13.439 | 0.617 | 0.977 | 0.000 | 50 | 0.617 | -0.007 | 13.209 | 0.990 | 0.000 | 85 | 4554045 | 50 |
|  | IBD | Healthy | 0.376 | 14.110 | 0.702 | 0.976 | 0.000 | 50 | 0.525 | -0.007 | 13.879 | 0.994 | 0.000 | 72 | 5951880 | 50 |
|  |  | Disease | 0.392 | 14.013 | 0.688 | 0.978 | 0.000 | 50 | 0.543 | -0.007 | 13.781 | 0.995 | 0.000 | 74 | 5792650 | 50 |
|  | Mean of HEA |  | 0.390 | 13.946 | 0.689 | 0.977 | 0.000 | 50 | 0.534 | -0.006 | 13.709 | 0.994 | 0.000 | 84 | 5709650 | 50 |
|  | SE of HEA |  | 0.023 | 0.224 | 0.021 | 0.002 | 0.000 | 0 | 0.023 | 0.001 | 0.216 | 0.000 | 0.000 | 7 | 750330 | 0 |
|  | Mean of DIS |  | 0.409 | 13.866 | 0.672 | 0.977 | 0.000 | 50 | 0.556 | -0.007 | 13.627 | 0.993 | 0.000 | 85 | 5658598 | 50 |
|  | SE of DIS |  | 0.030 | 0.217 | 0.028 | 0.001 | 0.000 | 0 | 0.032 | 0.001 | 0.211 | 0.001 | 0.000 | 7 | 602755 | 0 |
| $q=1$ | Obesity | Lean | 0.325 | 12.432 | 0.748 | 0.954 | 0.000 | 50 | 0.474 | -0.006 | 12.165 | 0.981 | 0.000 | 85 | 980004.2 | 50 |
|  |  | Overweight | 0.301 | 12.375 | 0.768 | 0.930 | 0.000 | 50 | 0.478 | -0.007 | 12.057 | 0.972 | 0.000 | 73 | 831126.0 | 49 |
|  | Type-II diabetes | Healthy | 0.321 | 11.721 | 0.751 | 0.941 | 0.000 | 50 | 0.489 | -0.008 | 11.461 | 0.973 | 0.000 | 61 | 435659.0 | 49 |
|  |  | Disease | 0.363 | 11.679 | 0.714 | 0.940 | 0.000 | 50 | 0.534 | -0.008 | 11.418 | 0.966 | 0.000 | 64 | 491949.4 | 46 |
|  | IBD | Healthy | 0.264 | 12.514 | 0.799 | 0.920 | 0.000 | 50 | 0.433 | -0.008 | 12.251 | 0.966 | 0.000 | 53 | 757500.7 | 48 |
|  |  | Disease | 0.337 | 12.316 | 0.736 | 0.947 | 0.000 | 50 | 0.534 | -0.009 | 12.007 | 0.982 | 0.000 | 57 | 831436.9 | 47 |
|  | Mean of HEA |  | 0.303 | 12.222 | 0.766 | 0.938 | 0.000 | 50 | 0.465 | -0.007 | 11.959 | 0.974 | 0.000 | 66 | 724388.0 | 49 |
|  | SE of HEA |  | 0.020 | 0.252 | 0.017 | 0.010 | 0.000 | 0 | 0.017 | 0.001 | 0.250 | 0.004 | 0.000 | 9 | 158008.7 | 1 |
|  | Mean of DIS |  | 0.334 | 12.123 | 0.739 | 0.939 | 0.000 | 50 | 0.516 | -0.008 | 11.827 | 0.973 | 0.000 | 65 | 718170.8 | 47 |
|  | SE of DIS |  | 0.018 | 0.223 | 0.016 | 0.005 | 0.000 | 0 | 0.019 | 0.001 | 0.205 | 0.005 | 0.000 | 5 | 113110.7 | 1 |
| $q=2$ | Obesity | Lean | 0.325 | 11.061 | 0.747 | 0.908 | 0.000 | 50 | 0.527 | -0.007 | 10.699 | 0.956 | 0.000 | 71 | 247026.3 | 47 |
|  |  | Overweight | 0.265 | 11.063 | 0.798 | 0.811 | 0.000 | 50 | 0.501 | -0.009 | 10.641 | 0.903 | 0.000 | 58 | 193637.8 | 45 |
|  | Type-II diabetes | Healthy | 0.244 | 10.549 | 0.816 | 0.856 | 0.000 | 50 | 0.437 | -0.009 | 10.248 | 0.922 | 0.000 | 49 | 99610.3 | 46 |
|  |  | Disease | 0.317 | 10.408 | 0.755 | 0.871 | 0.000 | 50 | 0.528 | -0.010 | 10.096 | 0.916 | 0.000 | 51 | 113517.0 | 44 |
|  | IBD | Healthy | 0.221 | 11.266 | 0.835 | 0.820 | 0.000 | 50 | 0.437 | -0.010 | 10.936 | 0.914 | 0.000 | 42 | 186203.9 | 44 |
|  |  | Disease | 0.334 | 10.950 | 0.740 | 0.890 | 0.000 | 50 | 0.579 | -0.012 | 10.563 | 0.954 | 0.000 | 50 | 208940.3 | 45 |
|  | Mean of HEA |  | 0.263 | 10.958 | 0.799 | 0.862 | 0.000 | 50 | 0.467 | -0.009 | 10.628 | 0.931 | 0.000 | 54 | 177613.5 | 46 |
|  | SE of HEA |  | 0.032 | 0.213 | 0.027 | 0.026 | 0.000 | 0 | 0.030 | 0.001 | 0.202 | 0.013 | 0.000 | 9 | 42771.5 | 1 |
|  | Mean of DIS |  | 0.305 | 10.807 | 0.764 | 0.858 | 0.000 | 50 | 0.536 | -0.010 | 10.433 | 0.924 | 0.000 | 53 | 172031.7 | 45 |
|  | SE of DIS |  | 0.021 | 0.202 | 0.018 | 0.024 | 0.000 | 0 | 0.023 | 0.001 | 0.170 | 0.015 | 0.000 | 3 | 29588.9 | 0 |
| $q=3$ | Obesity | Lean | 0.346 | 9.945 | 0.729 | 0.879 | 0.000 | 50 | 0.561 | -0.008 | 9.552 | 0.925 | 0.000 | 72 | 88601.1 | 46 |
|  |  | Overweight | 0.273 | 9.934 | 0.791 | 0.760 | 0.000 | 48 | 0.487 | -0.008 | 9.578 | 0.847 | 0.000 | 59 | 64918.8 | 43 |
|  | Type-II diabetes | Healthy | 0.220 | 9.669 | 0.835 | 0.804 | 0.000 | 50 | 0.404 | -0.009 | 9.379 | 0.883 | 0.000 | 48 | 37648.9 | 46 |
|  |  | Disease | 0.319 | 9.350 | 0.752 | 0.809 | 0.001 | 49 | 0.505 | -0.010 | 9.111 | 0.840 | 0.001 | 52 | 40409.1 | 39 |
|  | IBD | Healthy | 0.190 | 10.088 | 0.859 | 0.729 | 0.001 | 46 | 0.382 | -0.010 | 9.831 | 0.825 | 0.000 | 39 | 51321.6 | 41 |
|  |  | Disease | 0.294 | 9.676 | 0.774 | 0.760 | 0.001 | 46 | 0.523 | -0.012 | 9.369 | 0.827 | 0.000 | 45 | 50731.4 | 41 |
|  | Mean of HEA |  | 0.252 | 9.900 | 0.808 | 0.804 | 0.000 | 49 | 0.449 | -0.009 | 9.587 | 0.878 | 0.000 | 53 | 59190.5 | 44 |
|  | SE of HEA |  | 0.048 | 0.123 | 0.040 | 0.043 | 0.000 | 1 | 0.056 | 0.001 | 0.132 | 0.029 | 0.000 | 10 | 15225.8 | 2 |
|  | Mean of DIS |  | 0.295 | 9.653 | 0.773 | 0.776 | 0.001 | 48 | 0.505 | -0.010 | 9.353 | 0.838 | 0.000 | 52 | 52019.8 | 41 |
|  | SE of DIS |  | 0.013 | 0.169 | 0.011 | 0.016 | 0.000 | 1 | 0.010 | 0.001 | 0.135 | 0.006 | 0.000 | 4 | 7104.6 | 1 |

\* $N$ =The number of times (number of random re-samplings) that the $m$ -DAR model was successfully fitted.

#### MAD (maximal accrual diversity of metagenome) with DAR-PLEC Models

Ma (2018a) derived the maximal accrual diversity (MAD) in a cohort or population based on the DAR-PLEC model (eqn. 3) as follows:

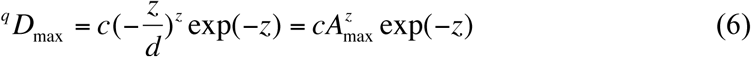

and the number of individuals (*A*_max_) reaching the maximum (*D*_*max*_) can be estimated by

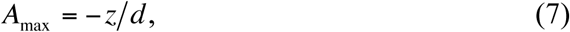

where all parameters are the same as eqns. (3) & (5).

Similar to the previous definition for *m*-DAR profile (*z-q* pattern), we define *m*-MAD profile (*D*_max_-*q* pattern) as a series of *D*_max_ values corresponding to different diversity order (*q*). This parameter did not exist in classic SAR but can be harnessed to measure the diversity accrual in a population or cohort setting (Ma 2018a).

MAD (*D*_max_) can act as a proxy for the so-termed potential (‘dark’) diversity, which refers to the contribution of species that are absent locally but present in regional species poor on regional or global scales (Partel *et al*. 2011, Real *et al*. 2017, Ma 2019). Since the diversity order (*q*) of Hill numbers represents different weights by species (gene) abundance distributions, and the Hill numbers at different orders essentially are the number of species (genes) with different level of rarity/commonness. Therefore, MAD (*D*_max_) can be used to estimate the potential diversity on the whole spectrum of rarity *vs*. commonness, *e.g*., the potential diversity of common genes *vs*. dominant genes.

#### PDO (pair-wise diversity overlap) profile

The *pair-wise diversity overlap* (*g*) of two bordering areas of the same size (*i.e*., the proportion of the new diversity in the second area) is:

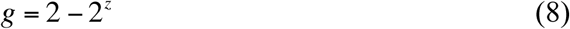

where *z* is the scaling parameter of *m*-DAR PL model [eqns. (2) & (4)]. When *z*=1, then *g*=0, no overlap exists; and when *z*=0, *g*=1, total overlap exists. In reality, *g* should be between 0 and 1.

Since the *equal size* of *area* assumption is largely true in the case of sampling human microbiome, the parameter z of the PL-DAR can be utilized to estimate the *pair-wise diversity overlap* (PDO), *i.e*., the diversity overlap between two individuals in the human microbiome with Eqn. (8). Given the range of *g* is between 0 and 1, we may even use percentage (%) notation to measure pair-wise diversity overlap.

Similar to previous definitions for *m*-DAR profile (*z-q* pattern) and *m*-MAD profile (*D*_max_-*q* pattern), the *m*-PDO profile (*g-q* pattern) is defined as a series of *g*-values of the pair-wise diversity overlap metric (*g*) at different diversity order (*q*). It approximates the similarity between a pair of individuals in the case of human metagenomes (in which the size of ‘area’ is approximately same).

#### RIP (Ratio of individual- to population-diversity) Profile

We define the RIP (ratio of Individual diversity to Population accrual diversity) as:

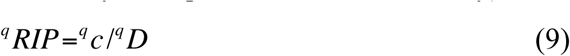

where *c* is the *m*-DAR-PL parameter, and ^*q*^*D* is the estimated accrual diversity of the population (cohort), both at diversity order *q* and computed with eqn. (2). We further define ^*q*^*RIP-q* series (there is a *RIP* for each diversity order *q*) as RIP profile, similar to the previously defined DAR-, PDO-, and MAD-profiles.

According to the above RIP definition, a RIP profile can be constructed with population (cohort) of any size, however, in practice, using ^*q*^*D*_*max*_ in place of ^*q*^*D* should be more convenient to apply, that is:

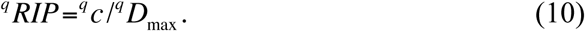

The RIP parameter measures the average level of an individual can represent a population (or cohort) from which the individual comes from. As argued previously, parameter *c* is an *approximated* value of individual diversity (diversity per individual). The approximation is contingent on two implicit assumptions: (*i*) the sizes of areas are equal, which is generally true in the case of human microbiome; (*ii*) the start of area accrual won’t exert significant influence on the estimation of parameter *c*. This appears to be satisfied given that assumption (*i*) is largely true for the human microbiome. However, it is well known that human microbiome is highly heterogeneous or so-termed personalized. So the choice of starting area to accrue may indeed matter in estimating the parameter *c*. To deal with the issue associated with assumption (*ii*), we take the parameter average from 50 (or 100) times of repeated building of DAR-PLEC models for a population (cohort) as explained previously. Without this repetition, parameter *c* cannot represent the *average* individual diversity because of the high heterogeneity among individuals in their metagenome diversities. Therefore, the usage of RIP should be cautioned in practice for the reasons explained above. Occasionally, the datasets may not allow for reliable estimation of the MAD (e.g., PLEC model fails to fit data), we suggest using the predicted value from PL model itself, *i.e*., using Eqn. (9).

## Results

### *m*-DAR (metagenome-DAR) for the MG (metagenomic gene) diversity

#### PL (power law) models for the *m*-DAR of MG diversity

From the PL model (the left side in Table 2) for the *m*-DAR in terms of the MG diversity of the human gut metagenome, we summarize the following findings:

i. The *m*-DAR PL model fitted to all six datasets of the gut metagenome, including 6 treatments of 3 healthy and 3 diseased treatments (obesity, diabetes and IBD), extremely significant (*p-* value<0.001).
ii. The ranges of *m*-DAR PL scaling parameter (*z*) for the MG (metagenomic gene) were [0.358, 0.468] at diversity order *q*=0, [0.264, 0.363] at *q*=1, [0.221, 0.334] at *q*=2, and [0.190, 0.346] at *q*=3, respectively. These ranges are rather similar to the species-DAR for the gut microbiome reported in Ma (2018a), which are rather close the counterpart of species-DAR at the diversity order *q*=0 [0.293, 0.332], but rather divergent from the counterparts of species-DAR when *q*>0, with *s*-DAR scaling parameter (*z*) being significantly small (*z<*0.1). That is, the scaling rates (*z*) of *gene richness* and *species richness* (the carriers of those genes) are rather close to each other, but the scaling rates of general diversity (*q*>0) are rather different between species diversity and gene diversity, with MG scaling (*z*) having a much larger magnitude.
iii. Table 2 also listed the comparative results between the healthy and diseased treatments, including the parameters for each treatment, as well as the average parameters for the healthy *vs*. diseased treatments across all three case studies. Overall, the differences between the healthy and diseased treatments in their *m*-DAR parameters are relative small. For example, the average difference in the scaling parameter (*z*) between the healthy and diseased treatments was 0.019 for *q*=0, 0.031 for *q*=1, 0.042 for *q*=2; and 0.043 for *q*=3, respectively, with the diseased treatments being slightly higher in all diversity orders. As shown in Table S1, the randomization test indicated that there was no significant difference between the healthy and diseased treatments in all three case studies (*p*>0.05). In other words, the three diseases (obesity, diabetes, and IBD) do not significantly influence the scaling of MG diversity.
iv. The pattern of PDO (pair-wise diversity overlap) profile (*i.e*., *g-q* series) exhibited the opposite pattern as the DAR profile (*z*-*q*) did, as expected. This is because, while the scaling parameter *z* (DAR-profile) measures the heterogeneity of neighboring areas, the PDO-profile parameter (*g*) measures the homogeneity (overlap or similarity) of neighboring areas.
v. The parameter ln(*c*) ranged [9.350, 14.224] across all diversity orders, corresponding a range of *c* [10^4^, 1.5×10^6^] approximately. The values of *c* are magnitudes larger than those of microbial species reported in Ma (2018a, 2018b). This is expected given that the *gene richness* and *diversity* are magnitudes higher than *species richness* and *diversity*. Parameter *c* is a rough estimate of the metagenomic genes (MG) carried by one individual (when *A*=1).

#### PLEC (power law with exponential cutoff) models for the *m*-DAR of MG diversity

From the PLEC model (the right side in Table 2) of the *m*-DAR of the MG (metagenomic gene) diversity of the human gut metagenome, we summarize the following findings:

i. The PLEC model fitted to all six datasets extremely significant with *p*-value<0.001, actually fitted slightly better than the PL model as indicated by the higher *R* (linear correlation coefficient) column.
ii. The negative value of parameter *d* of the PLEC model showed that there was an asymptotic line, which was represented by *D*_*max*_ and corresponding *area* size (*i.e*., *A*_*max*_ or the number of individual subjects accumulated to reach the asymptote). Indeed, the most important contribution the PLEC model brings in is the estimation of the MAD (*D*_*max*_: maximal accrual diversity) of MGs. To the best of our knowledge, this should be the first estimation of the gene diversity accrual in a population setting. For example, at *q*=0, the estimated MAD of MGs ranged between [4,305,968, 6,871,100] and its standard error ranged [602,755, 750,330]. The range of MAD of MGs, which can be considered as a measure of the *gene richness at the population level* (or *gene richness accrual*) is one magnitude higher than the *gene richness of an individual* exhibited in Table 1, which ranged [504,712, 973,838]. That is, the range of the standard error of MAD of MGs is actually close to the gene richness of an individual (see Table 1). This suggests that the magnitude of *variation* of MAD within a population is close to the total number of metagenomic gene species (kinds) from an individual carries, an exceptionally high inter-subject heterogeneity.

Fig 2 & 3 displayed the graphs for the scaling parameter (*z*) and maximal accrual diversity (*D*_*max*_) of the MG for the six treatments of three case studies at different diversity orders (*q*=0-3).

**Fig 2.**
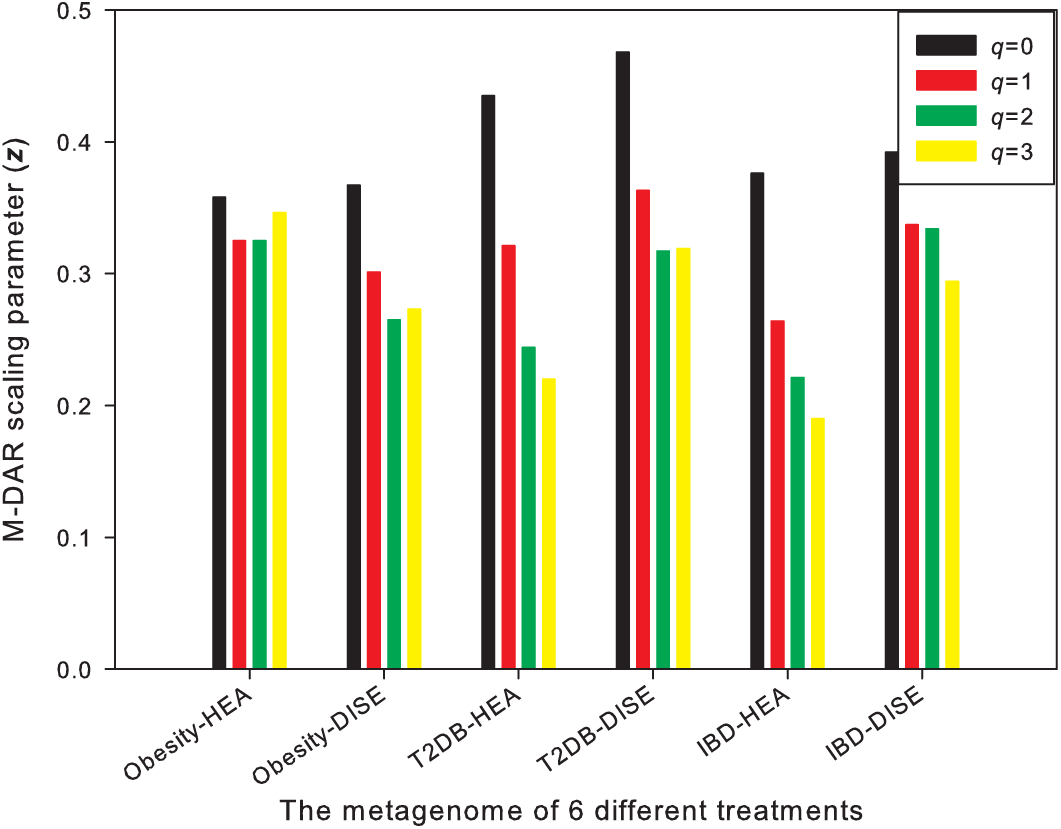
The scaling parameter (*z*) of the *m*-DAR (metagenome-diversity area relationship) for the metagenomic-genes (MGs) of the human gut metagenome

**Fig 3.**
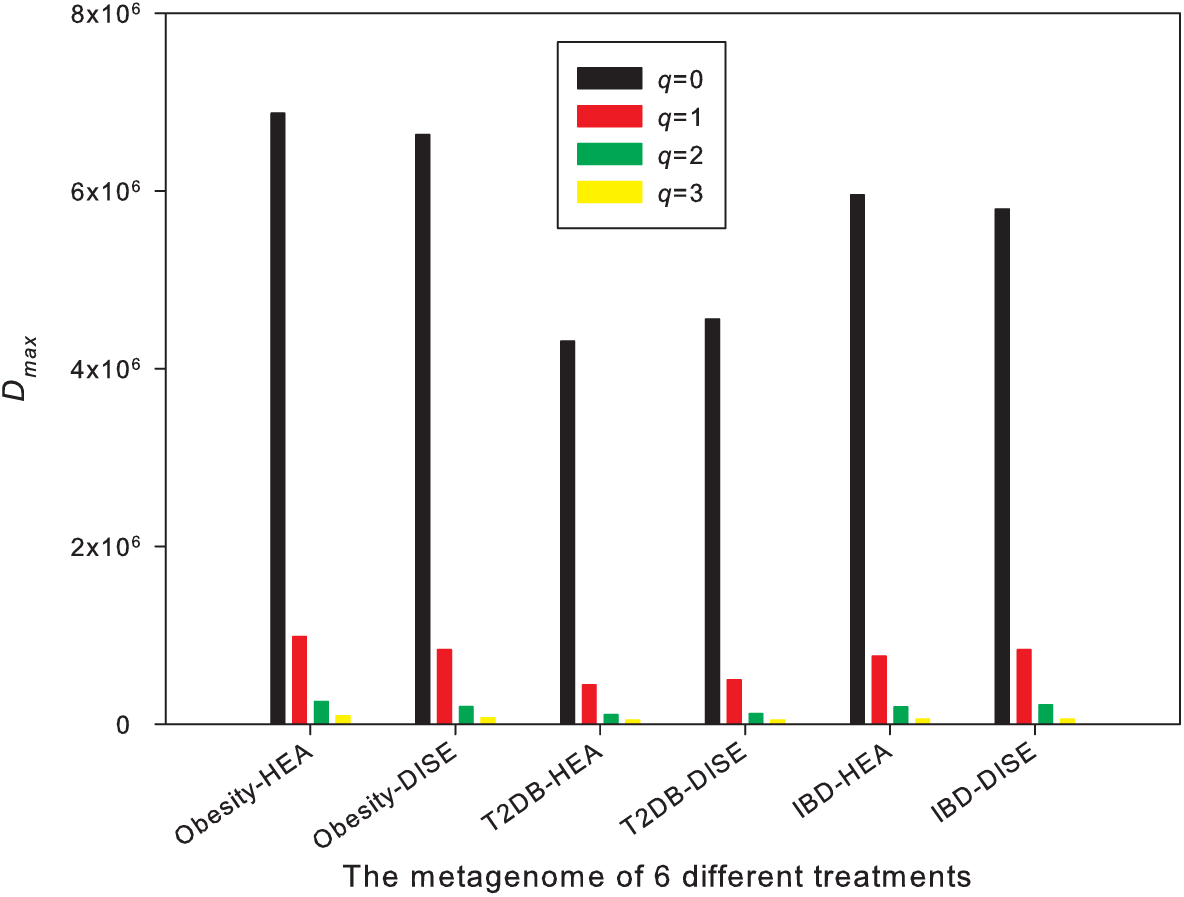
The MAD (maximum accrual diversity: *D*_*max*_) of the *m*-DAR (metagenome diversity-area relationship) for the MG (metagenomic genes) of the human gut metagenome

### *m*-DAR for the MFGC (metagenome functional gene cluster) diversity

#### PL models for the *m*-DAR of MFGC diversity

We further built *m*-DAR PL models with the MFGC abundance table (the left side of Table 3). The difference between the *m*-DAR models built here and the *m*-DAR models for the MG in the previous section is that, here the metagenomic entity to be accumulated is MFGC (metagenome functional gene cluster), rather than the basic metagenomic gene (MG) as in the previous section.

**Table 3.**
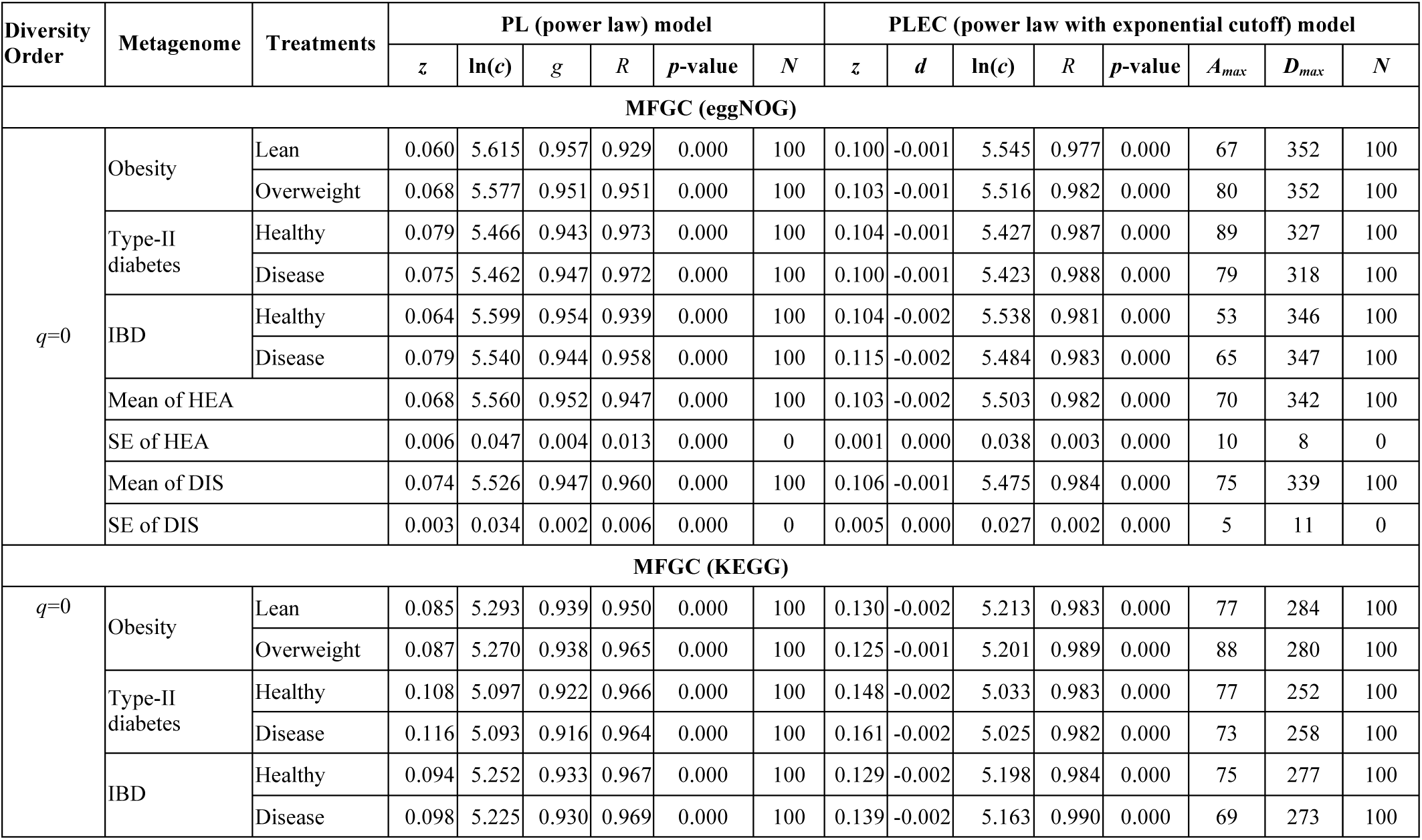
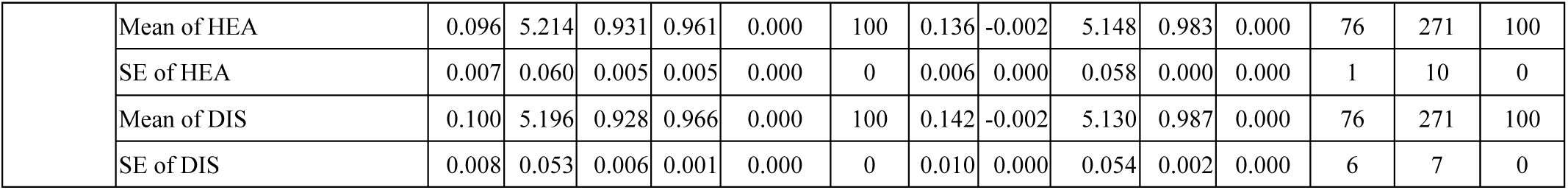
The parameters of m-DAR (metagenome diversity-area relationship) models fitted for MFGC (metagenome functional gene cluster) diversity, averaged from 100 times of re-sampling

From the PL model of m-DAR (the left side in Table 3) for the MFGC diversity scaling across individuals of a cohort (or population), we summarize the following findings:

i. The *m*-DAR PL model was successfully fitted to the MFGC diversity in all 100 times of re-sampling (100% of success) when *q*=0, but only succeeded in approximately 80% of the re-sampling when *q*>0. Considering that a MFGC is a functional unit *cluster* of MGs, MFGCs should be much more homogenous than the MGs among individuals, and the number of MFGCs (*i.e*., MFGC *richness*) should already be rather informative as a diversity measure. Hence, there is no need to use more complex high-order Hill numbers. Therefore, we suggest to limit the *m*-DAR modeling for MFGC to the diversity order of *q*=0, *i.e*., the *richness* of MFGC.
ii. The *m*-DAR PL scaling parameter (*z*) for the MFGC is less than 1/5 of that for the MG, which supports our previous statement that MFGCs are much more homogenous than MGs among individual subjects, because obviously smaller scaling parameter (*z*) indicates slower changes (*i.e*., high homogeneity or similarity) in diversity accrual with individuals (areas).
iii. Regarding the MFGC clustered based on the eggNOG database, the mean scaling parameter (*z*) of the healthy treatments is about 0.068, and the mean *z* of the diseased treatments is about 0.074, with the diseased being slightly higher. Regarding the MFGC clustered based on the KEGG database, the mean *z* of the healthy treatments is about 0.096, and the mean *z* of the diseased treatments is about 0.100, with the diseased being slightly higher too. Statistically, the randomization test indicates that there was no significant difference between the healthy and diseased treatments in all three case studies (*p*>0.05) (Table S2).
iv. The pair-wise diversity overlap (PDO or *g*) for the MFGC is much larger than the PDO for the MG. The high PDO in MFGC simply confirmed the high mutual similarity (overlap) or lower heterogeneity among individuals in terms of MFGC abundance distribution than in terms of the MG, as revealed in (*ii*) & (*iii*).

#### PLEC models for the *m*-DAR of the MFGC diversity

From the PLEC model of the *m*-DAR (the right side in Table 3) for MFGC diversity scaling across individuals within a cohort (or population), we summarize the following findings:

i. With the PLEC model for MFGC, we can predict the MAD (maximal accrual diversity) of MFGC. Given that we limit the diversity order for MFGC to the *q*=*0*-th order as explained previously, the term “*maximum accrual richness*” (MAR) of MFGC should be a term that more accurately describes its metagenomic meanings. To the best of our knowledge, there was not an approach to estimating this number prior to our work. Note that the MAR of MFGC (in a cohort or population) is rather close to the average MFGC of an individual metagenome contains. This again confirmed the lower heterogeneity (or high homogeneity) among individuals in their MFGC richness.
ii. The difference between the healthy and diseased treatments in *D*_max,_ *i.e*., the MAR of MFGC is statistically insignificant (Table S2, *p*>0.05). The difference was a loss of 3 MFGCs in the diseased metagenomes for the eggNOG-based MFGC, and 0 loss (*i.e*., the exactly same number) for the KEGG-based MFGC. This is, of course, as expected, because losing a functional gene cluster such as a pathway in the case of KEGG could be deadly for the survival of microbes, and most disturbances including disease usually do not cause such lethal effect.

Fig 4 & 5 displayed the key parameters of the *m*-DAR models for MFGC diversity, including diversity scaling parameter (*z*), pair-wise diversity overlap (similarity) parameter (*g*), and MAR (*D*_*max*_).

**Fig 4.**
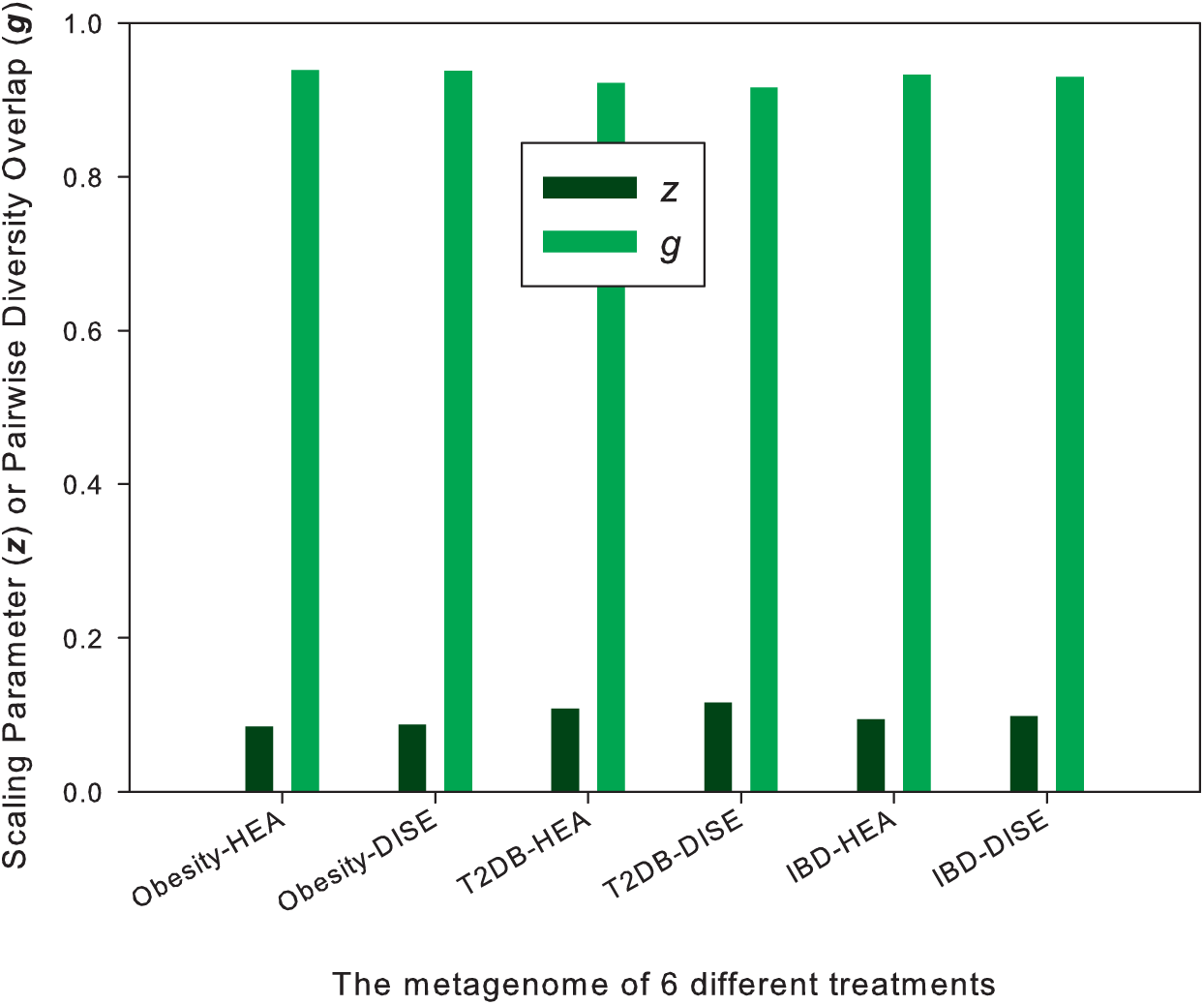
The scaling parameter (*z*) and PDO (pair-wise diversity overlap) parameter (*g*) of the *m*-DAR (metagenome diversity-area relationship) models for the MFGC (metagenome functional gene cluster) diversity, with 3 pairs of the healthy (HEA) and diseased (DISE) treatments, from 3 case studies (obesity, type-II diabetes, and IBD) (in the case of MFGC, we build *m*-DAR models for *q*=0 only)

**Fig 5.**
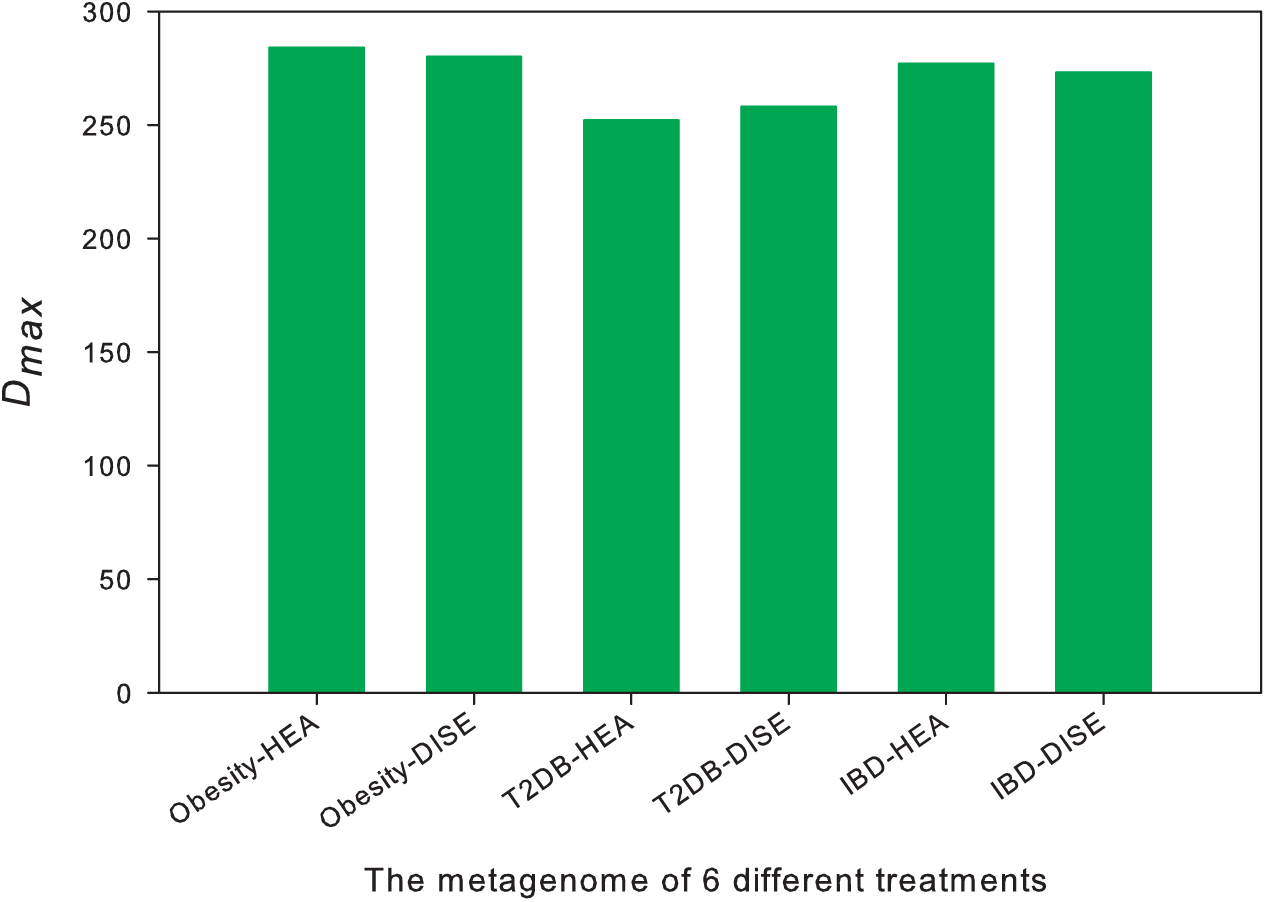
The MAD (maximum accrual diversity) of the *m*-DAR (metagenome-diversity area relationship) for the MFGC (metagenome functional gene cluster), with 3 pairs of the healthy (HEA) and diseased (DISE) treatments, from 3 case studies (obesity, type-II diabetes, and IBD) (In the case of MFGC, we build m-DAR models for *q*=0 only)

### The RIP (Ratio of Individual diversity to Population accrual diversity)

According to the definition of PIP (Eqn. 9), it is the ratio (percentage) of an average individual diversity (the parameter *c* of PL model) to the accrued diversity of the population (or cohort). In most cases, we suggest to use the MAD (maximum accrued diversity) or *D*_*max*_, which is the theoretical maximum or asymptote line of *m*-DAR PLEC model, as a replacement for population accrual diversity (*i.e*., using Eqn. 10, instead).

Table 4 listed the estimated PIP profile in terms of the MG and MFGC, for the healthy and diseased treatments, respectively. In the case of MFGC, only RIP for *q*=0 is computed since we suggested to limit the *m*-DAR modeling for MFGC abundance tables (data) to *q*=0, or MFGC richness. Three findings can be discovered from Table 4:

i. In terms of the MG, an average individual (person) may carry approximately 1/5-1/3 (the range or variation depends on *q*) of the MGs carried by the whole population, while in terms of the MFGC, the ratio is approximately 2/3. This should be expected because MFGCs are clusters of MGs and contain lots of functionally redundant MGs.
ii. The average RIP for the healthy treatment is slightly higher than that of the diseased treatment, but the difference may not be statistically significant.
iii. The RIP exhibited lower percentages at the lower-diversity orders than at the high-diversity orders. That is, at higher diversity orders, the heterogeneity in diversity becomes smaller, and an individual is more representative of the population he or she belongs to.

**Table 4.** The RIP (the ratio of individual diversity to population MAD), averaged from the RIP of the three case studies on the human gut metagenome (converted to percentage)

| Diversity Order ($q$) | MG | | MFGC | |
| --- | --- | --- | --- | --- |
|  | Average RIP (%) For Healthy | Average RIP (%) For Diseased | Average RIP (%) For Healthy | Average RIP (%) For Diseased |
| $q=0$ | 20.0 | 18.6 | 67.8 | 66.6 |
| $q=1$ | 28.1 | 25.6 | N/A | |
| $q=2$ | 32.3 | 28.7 | | |
| $q=3$ | 33.7 | 29.9 | | |

## Conclusions and Discussion

We introduced the *m*-DAR models (both PL and PLEC) for investigating the scaling of MG and MGFC, and defined four biogeography profiles (including DAR-profile, PDO profile, MAD profile and RIP profile) for characterizing the biogeography of microbial metagenome based on the *m*-DAR parameters. We demonstrated the applications of those models and profiles with the metagenomic datasets of human gut microbiomes. For example, the *m*-MAD profile provides a valuable tool to estimate the *potential* MG or MFGC diversity of a human population. According to the estimation from *m*-MAD profile (Table 2), the *potential* total number of MGs is up to 6.8 million (*q*=0). Among the total number of MGs, commonly encountered MGs are about 1/7 of the total MGs or 1 million (*q*=1), the dominant (more abundant than the common) MGs are about 3% or 200,000 (*q*=2), and most dominant (much more abundant) MGs are about 1% or 60,000 of the total MGs.

The current focusing on the distribution of the diversity of organisms, while ignoring the *diversity* of genomes or metagenomes, has at least two main reasons. While the primary tool (morphology-based taxonomy) for investigating traditional (systematic) biogeography has been available perhaps since ancient time, at least, since 18^th^ century of Linnaean taxonomy, the primary tool for identifying genome/metagenome, DNA sequencing technology, is a very recent technology, only readily accessible in the last two decades. Besides the delayed availability of the technology for measuring gene/metagenome diversity (for investigating evolutionary biogeography), another reason has been the missing of a theory and supporting tool to utilize the metagenome diversity information and to add a new *metagenomic dimension* to existing biogeography of organisms. As rightly indicated by Boon *et al*. (2013) “A central challenge in microbial community ecology is the delineation of appropriate units of biodiversity, which can be taxonomic, phylogenetic, or functional in nature.” The *m*-DAR we develop in this study offers such an example of the theoretical model and tool. Integrated together, both *s*-DAR and *m*-DAR demonstrated the feasibility of a unifying approach to bridging possible gaps between systematic biogeography and evolutionary biogeography, also of an inclusive biogeography of the biomes of plants, animals and microbes.

Previously, Zhou *et al*. (2008) conducted a pioneering application of the traditional SAR to the metagenomic functional genes in forest soil microbiome. Perhaps restricted by then available sequencing technology a decade ago, their study was limited to functional gene clusters without measuring the metagenomic genes, which rely on high throughput NGS sequencing technologies. The range of scaling parameter (*z*) they obtained was 0.048-0.096, which is rather close to the range we obtained for the MFGC of the human gut microbiome (Table 3) in this study (0.060-0.116). To the best of our knowledge, there are not any other comparable results with our *m*-DAR parameters for the MG in existing literature. Hence, no further comparative observations were noted in this study.

As a side note, although we could potentially transform the DTR (diversity-time period relationship) (Ma 2018c) into metagenome-DTR, we failed to locate long enough time-series datasets of metagenomic whole-genome sequencing data. This is obviously due to the still relatively high cost of whole-genome sequencing, which is currently approximately 10 times of the amplicon-sequencing cost of 16S-rRNA genes (used for species-DTR modeling). Indeed, it should be feasible to demonstrate the spatial-temporal changes of a human population with the metagenome version of DTAR (diversity-time-area relationship) (Ma 2018c, 2019) when future whole-genome sequencing costs drop to sufficiently low.

## Data accessibility

All six metagenomic datasets reanalyzed in this study are available in public domain and their sources are noted in Table 1.

## Author Contributions

ZS Ma designed and conducted the analysis, interpreted the results, and wrote the manuscript. AM Ellison interpreted the results and revised the manuscript. All authors approved the submission.

## Acknowledgements

This study received funding from the following sources: A National Natural Science Foundation (NSFC) Grant (No. 31970116) on “The medical ecology of human microbiome associated diseases”; the Cloud-Ridge Biotech Industry Leader award; A China-US Collaborative Project on Genomics/Metagenomics Big Data.

## List of Online Supplementary Tables

**Table S1.**
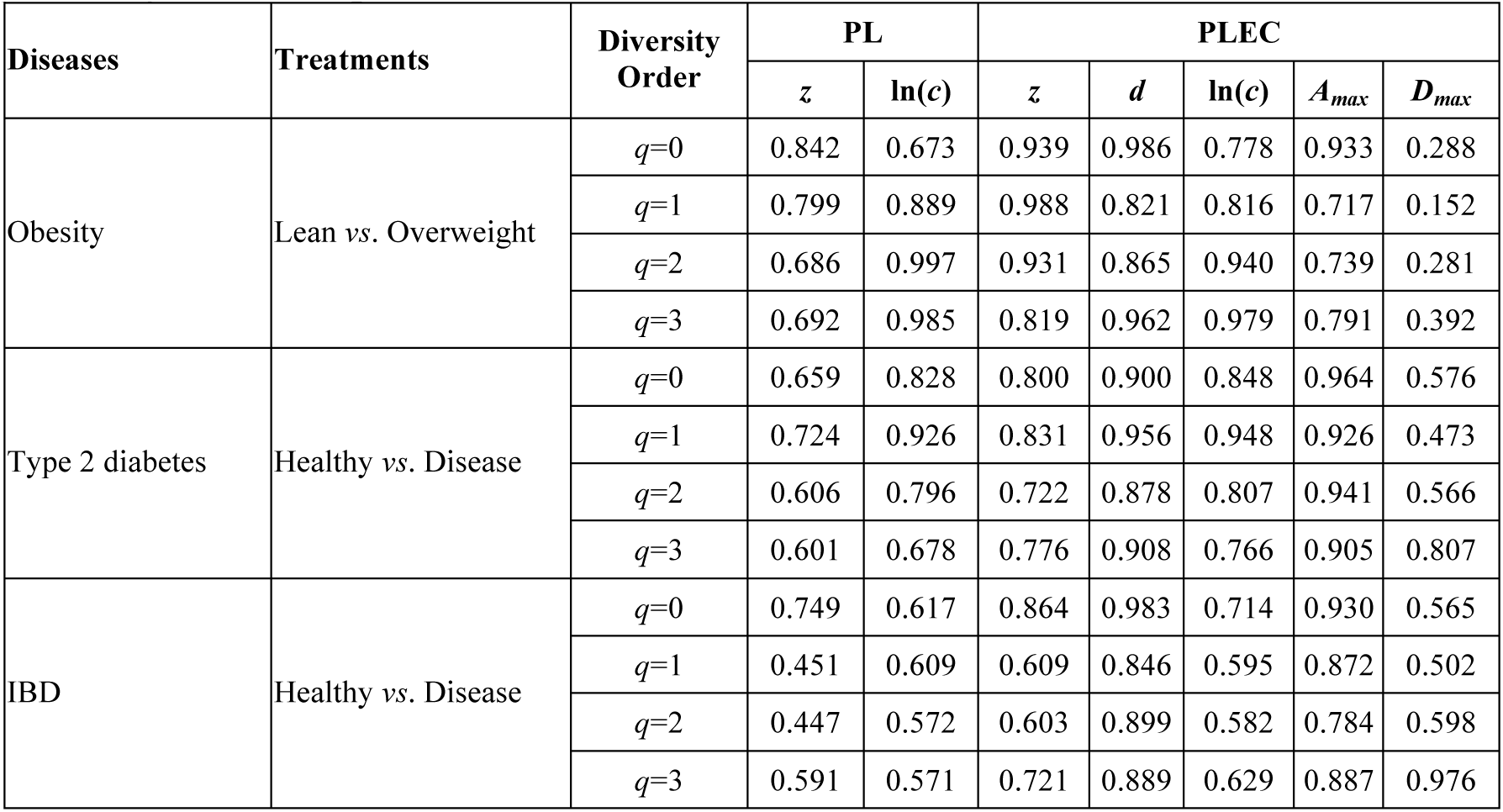
The *p*-value of the randomization test for the difference in the *m*-DAR parameters of the MGs (metagenomic genes) between the healthy and diseased treatments of the human gut metagenome samples

**Table S2.** The *p*-value of the randomization test for the difference in the *m*-DAR parameters of the MFGCs (metagenome functional gene clusters) between the healthy and diseased treatments of the human gut metagenome samples

| Microbiome | Treatments | Diversity Order | PL |  | PLEC |  |  |  |  |
| --- | --- | --- | --- | --- | --- | --- | --- | --- | --- |
| | | | $z$ | $\ln(c)$ | $z$ | $d$ | $\ln(c)$ | $A_{max}$ | $D_{max}$ |
| MFGC Type-I (eggNOG) |  |  |  |  |  |  |  |  |  |
| Obesity | Lean vs. Overweight | $q=0$ | 0.575 | 0.512 | 0.918 | 0.772 | 0.721 | 0.489 | 0.868 |
| Type 2 diabetes | Healthy vs. Disease | $q=0$ | 0.748 | 0.956 | 0.888 | 0.949 | 0.953 | 0.922 | 0.637 |
| IBD | Healthy vs. Disease | $q=0$ | 0.423 | 0.402 | 0.738 | 0.851 | 0.542 | 0.549 | 0.906 |
| MFGC Type-I (KEGG) |  |  |  |  |  |  |  |  |  |
| Obesity | Lean vs. Overweight | $q=0$ | 0.885 | 0.649 | 0.862 | 0.732 | 0.878 | 0.658 | 0.497 |
| Type 2 diabetes | Healthy vs. Disease | $q=0$ | 0.729 | 0.967 | 0.787 | 0.863 | 0.957 | 0.915 | 0.549 |
| IBD | Healthy vs. Disease | $q=0$ | 0.825 | 0.695 | 0.798 | 0.819 | 0.697 | 0.822 | 0.571 |
| MFGC Type-II (eggNOG) |  |  |  |  |  |  |  |  |  |
| Obesity | Lean vs. Overweight | $q=0$ | 0.476 | 0.429 | 0.734 | 0.965 | 0.588 | 0.646 | 0.902 |
| Type 2 diabetes | Healthy vs. Disease | $q=0$ | 0.583 | 0.824 | 0.712 | 0.936 | 0.781 | 0.967 | 0.683 |
| IBD | Healthy vs. Disease | $q=0$ | 0.531 | 0.524 | 0.850 | 0.822 | 0.667 | 0.554 | 0.862 |
| MFGC Type-II (KEGG) |  |  |  |  |  |  |  |  |  |
| Obesity | Lean vs. Overweight | $q=0$ | 0.826 | 0.629 | 1.000 | 0.852 | 0.770 | 0.774 | 0.454 |
| Type 2 diabetes | Healthy vs. Disease | $q=0$ | 0.718 | 0.947 | 0.777 | 0.825 | 0.919 | 0.882 | 0.539 |
| IBD | Healthy vs. Disease | $q=0$ | 0.895 | 0.798 | 0.758 | 0.712 | 0.735 | 0.697 | 0.569 |

## References

Arrhenius O. 1921. Species and area. Journal of Ecology, vol. 9, 95–99.

Bradley PD, Bakker HC, Rocha EPC et al (2019) Ultrafast search of all deposited bacterial and viral genomic data. Nat. Biotechnol. 37: 152–159.

Bolyen E, Rideout JR, Dillon MR et al (2019) Reproducible, interactive, scalable and extensible microbiome data science using QIIME 2. Nature Biotechnology 37: 852–857.

Boon E, CJ Meehan, C Whidden, DHJ Wong, MGI Langille & RG. Beiko (2013) Interactions in the microbiome: communities of organisms and communities of genes. FEMS Microbiol Rev 38 (2014) 90–118, DOI: 10.1111/1574-6976.12035

Chao A, Chiu CH, Hsieh TC. (2012). Proposing a resolution to debates on diversity partitioning. Ecology 93(9): 2037–2051

Chao A, CH Chiu, & L Jost (2014) Unifying species diversity, phylogenetic diversity, functional diversity and related similarity and differentiation measures through Hill numbers. Annual Reviews of Ecology, Evolution, and Systematics, 45, 297–324.

Chatelier EL, Nielsen T, Qin J, et al. (2013) Richness of human gut microbiome correlates with metabolic markers. Nature, 500:541–546.

Connor EF & ED McCoy (1979) The statistics and biology of the species–area relationship. The American Naturalist, 113, 791–833.

Costello EK, Stagaman K, Dethlefsen L, Bohannan BJM, Relman DA. (2012). The application of ecological theory toward an understanding of the human microbiome. Science 336:1255–1262.

Darwin, CR. (1859) On the origin of species London: John Murray. [1st edition]

Ebach, MC (2007) Introduction, In “Biogeography in a Changing World” ed. by Ebach, MC & RS Tangney (2007). CRC Press, Taylor & Francis Group, Florida, USA.

Forster SC, Kumar N, Anonye BO et al. (2019) A human gut bacterial genome and culture collection for improved metagenomic analyses. Nat. Biotechnol., 37: 186–192

Hanson, CA, JA. Fuhrman, M. Claire Horner-Devine and JBH Martiny (2012) Beyond biogeographic patterns: process shaping the microbial landscape. Nature Review Microbiology, 10:497–506.

Hill MO. (1973). Diversity and evenness: a unifying notation and its consequences. Ecology 54: 427–342.

Jost L. (2007). Partitioning diversity into independent alpha and beta components. Ecology 88: 2427–2439.

Li J, Jia H, Cai X, Zhong H, Feng Q, Sunagawa S et al. (2014). An integrated catalog of reference genes in the human gut microbiome. Nature Biotechnology 32(8): 834–841.

Lomolino MV. (2000). Ecology’s most general, yet protean pattern: the species–area relationship. Journal of Biogeography 27: 17–26.

Ma ZS, Li LW (2018) Measuring metagenome diversity and similarity with Hill numbers. Molecular Ecology Resources, 2018:1–17.

Ma ZS (2018a) Extending species-area relationships (SAR) to diversity-area relationships (DAR), Ecology and Evolution 8 (20): 10023–10038

Ma ZS (2018b) Sketching the Human Microbiome Biogeography with DAR (Diversity Area Relationship) Profiles. Microbial Ecology.

Ma ZS (2018c) Diversity time-period and diversity-time-area relationships exemplified by the human microbiome. Scientific Reports, 2018, 8(1):7214.

Ma ZS (2019) A new DTAR (diversity–time–area relationship) model demonstrated with the indoor microbiome. Journal of Biogeography, https://doi.org/10.1111/jbi.13636

Ma ZS, Ellison AM (2018) A unified concept of dominance applicable at both community and species scale. Ecosphere, https://doi.org/10.1002/ecs2.2477.

Ma ZS, Ellison AM (2019) Dominance network analysis provides a new framework for studying the diversity-stability relationship. Ecological Monographs. DOI: 10.1002/ecm.1358

Ma ZS, Li LW, Gotelli NJ (2019) Diversity-disease relationships and shared species analyses for human microbiome-associated diseases, The ISME Journal. DOI: 10.1038/s41396-019-0395-y

MacArthur, RH, Wilson, EO. (1967) The theory of islands biogeography. Princeton University Press. Princeton.

Martiny, JBH, J.M. Bohannan, JH. Brown, Robert K. Colwell, et al. (2006) Microbial biogeography: putting microorganisms on the map. Nature Review Microbiology, vol. 4, 102–112. doi: 10.1038/nrmicro1341

Nielsen HB, Almeida M, Juncker AS et al (2014) Identification and assembly of genomes and genetic elements in complex metagenomic samples without using reference genomes. Nature biotechnology, 32:822–828.

Plotkin JB et al. (2000). Predicting species diversity in tropical forests. Proceedings of the National Academy of Sciences USA 97: 10850–10854.

Partel, M., Szava-Kovats, R., & Zobel, M. (2011a). Dark diversity: Shedding light on absent species. Trends Ecology & Evolution., 26, 124–128. https://doi.org/10.1016/j.tree.2010.12.004

Preston FW (1960) Time and space and the variation of species. Ecology.

Qin J, Li R, Raes J et al. 2010. A human gut microbial gene catalogue established by metagenomic sequencing. Nature, 464:59–65.

Qin J, Li Y, Cai Z et al, 2012. A metagenome-wide association study of gut microbiota in type 2 diabetes. Nature, 490:55–60.

Real, R, Barbosa, A. M., & Bull, J. W. (2017). Species Distributions, quantum theory, and the enhancement of biodiversity measures. Systematic Biology, 66(3), 453–462.

Rosenzweig ML (1995) Species Diversity in Space and Time. Cambridge University Press, Cambridge.

Schloss PD, Westcott SL, Ryabin T, et al (2009) Introducing Mothur: Open Source, Platform-Independent, Community-Supported Software for Describing and Comparing Microbial Communities. Applied and Environmental Microbiology, 75, 7537–7541

Sczyrba A, Hofmann P, Belmann P, et al. (2017) Critical assessment of metagenome interpretation-a benchmark of metagenomics software. Nature Methods, 14(11):1063–1071.

Tjørve E. (2009). Shapes and functions of species–area curves (II): a review of new models and parameterizations. Journal of Biogeography 36: 1435–1445.

Ulrich W, Buszko J. (2003). Self-similarity and the species– area relation of Polish butterflies. Basic and Applied Ecology 4: 263–270.

van der Gast, C.J. (2013) Microbial biogeography and what Baas Becking should have said. Microbiology Today 40: 108–111.

van der Gast, CJ (2015) Microbial biogeography: the end of the ubiquitous dispersal hypothesis? Environmental Microbiology, 2015, 17, 3

Wang J & Jia H. (2016) Metagenome-wide association studies: fine-mining the microbiome. Nature Reviews Microbiology, 14(8):508–522.

Watson, HC. (1835) Remarks on geographic distribution of British plants. London.

Whittaker RJ, and KA Triantis (2012) The species–area relationship: an exploration of that ‘most general, yet protean pattern’. J of Biogeography, 39:623–626.

Xiao L, Feng Q, Liang S, et al. (2015) A catalog of the mouse gut metagenome. Nature Biotechnology, 33(10):1103–1108.

Xiao L, Estellé J, Kiilerich P, et al. (2016) A reference gene catalogue of the pig gut microbiome. Nature Microbiology, 1:16161.

Zhu W, Lomsadze A, Borodovsky M. 2010. Ab initio gene identification in metagenomic sequences. Nucleic Acids Res. 38, e132.

Zou Y, Xue W, Luo G et al. (2019) 1,520 reference genomes from cultivated human gut bacteria enable functional microbiome analyses. Nat. Biotechnol., 37: 179–185.

